# The Negev hospital-university-based (HUB) autism database

**DOI:** 10.1101/103770

**Authors:** Gal Meiri, Ilan Dinstein, Analya Michaelowski, Hagit Flusser, Michal Ilan, Michal Faroy, Asif Bar-Sinai, Liora Manelis, Dana Stolowicz, Lili Lea Yosef, Nadav Davidovitch, Hava Golan, Shosh Arbelle, Idan Menashe

**Affiliations:** Pre-School Psychiatry Unit, Soroka University Medical Center, Beer Sheva, Israel; Psychology Department, Ben-Gurion University of the Negev, Beer Sheva, Israel; Cognitive and Brain Sciences Department, Ben-Gurion University of the Negev, Beer Sheva, Israel; Zlotowski Center for Neuroscience, Ben-Gurion University of the Negev, Beer Sheva, Israel; Zusman Child Development Center, Soroka University Medical Center, Beer Sheva, Israel; Public Health Department, Ben-Gurion University of the Negev, Beer Sheva, Israel; Health Systems Management Department Ben-Gurion University of the Negev, Beer Sheva, Israel; Physiology and Cell Biology Department, Ben-Gurion University of the Negev, Beer Sheva, Israel

**Keywords:** Autism, Epidemiology, Multidisciplinary, Child development

## Abstract

Elucidating the heterogeneous etiologies of autism will require investment in comprehensive longitudinal data acquisition from large community based cohorts. With this in mind, we have established a hospital-university-based (HUB) database of autism which incorporates prospective and retrospective data from a large and ethnically diverse population. Here we present initial findings from 188 children who were diagnosed with autism during the first eighteen months of the study. The unique characteristics of this cohort included: significant differences between Bedouin and Jewish children in different risk factors and clinical characteristics; complete birth records for >90% of the children; and a high frequency of consanguineous marriages. Thus, the Negev HUB autism database comprises a remarkably unique resource to study different aspects of autism.

## Introduction

Autism is a heterogeneous neurodevelopmental disorder that is characterized by a wide spectrum of social and communication difficulties, repetitive and restricted behaviors, and a variety of clinical symptoms (Antshel et al. 2013; Georgiades et al. 2012; Kohane et al. 2012; Lai et al. 2013; Lenroot and Yeung 2013; Nazeer and Ghaziuddin 2012). While the DSM5 defined all individuals with autism as belonging to a single diagnostic category, it has been proposed that classification of autism into more homogenous subtypes will facilitate research, diagnosis, and treatment of the disorder (Dealberto 2013; Gabis and Pomeroy 2014; Kim et al. 2015; Lane et al. 2014). Previous classification schemes of autism that relied on behavioral and cognitive assessments (Association 2000; Levy et al. 2009; Lord and Jones) or restricted clinical measures (Kong et al. 2013; Lane et al. 2014; Lord and Jones; Ramsey et al. 2013) have failed to constitute a reliable basis for targeted treatment (Volkmar et al. 2012). Fortunately, the rapid progress in the fields of genomics, neuroimaging, and medical informatics provides exciting sources for new data that could be used for the development of potentially more useful classification approaches to autism.

In the last decade, research into the etiologies of autism is shifting from small-scale studies often focusing on one or several biological/clinical measures in a small sample, to more extensive studies that examine multiple types of data (e.g., genetic, neuroimaging, behavioral, and clinical) in larger cohorts (Buxbaum et al. 2014; Fischbach and Lord 2010; Geschwind et al. 2001; Matuszek and Talebizadeh 2009; Payakachat et al. 2015). This new approach has led to the identification of dozens of new susceptibility genes of autism (De Rubeis et al. 2014; Iossifov et al. 2012; Krumm et al. 2015; O’Roak et al. 2011; Sanders et al. 2012). Similar advances have been made in the identification of early neurophysiological (e.g., (Courchesne et al. 2011; Wolff et al. 2012)) and additional behavioral (e.g., (Chawarska and Shic 2009; Jones and Klin 2013)) characteristics associated with autism development. Nevertheless, many of the reported findings seem to vary substantially across different studies and populations, a factor that complicates generalization of conclusions (e.g., the existence of abnormally large head circumferences and its significance to early autism development (Zwaigenbaum et al. 2014)). Furthermore, many of these studies have relatively limited pre-diagnostic data, and usually do not collect sufficient follow-up data to “connect the etiological dots” from risk factors through biological mechanisms to precise phenotypic manifestations. There is, therefore, great motivation to construct autism databases containing both prospective and retrospective data that is relevant to multiple biological and clinical disciplines (e.g. genetics, metabolics, clinical history, behavioral assessments, neuroimaging, etc.) from the same individuals. Such information may enable in-depth characterization of specific autism etiologies and greatly improve our ability to classify autism into more homogeneous sub-types that may benefit from targeted treatments.

To address these issues, we have established a hospital-university-based (HUB) database of autism at the Soroka University Medical Center (SUMC) and Ben-Gurion University (BGU) in the Negev. SUMC is the only hospital in the Negev, which provides outpatient services to more than 75% of the Negev population (over 500,000 people), including child development and child psychiatry services. Our database was launched in January 2015 and already contains a wide variety of unique clinical and behavioral information regarding each participating child and his/her family members. These data include: behavioral and cognitive measures, socio-demographic data, clinical history, data on a wide verity of prenatal, perinatal, and neonatal risk factors, as well as information regarding sensory sensitivities and sleep quality. We are currently expanding our ongoing data collection to include additional behavioral and neuroimaging measures as well as genetic samples from this cohort, which is growing at a rate of ~16.5 new families per month. Here we present selected characteristics of ~300 families who were included in our database in the first 18 months of this initiative. The results demonstrate the remarkable potential of the Negev HUB database to be an extremely valuable resource for various interdisciplinary studies of autism.

## Methods

### Population

Most of the families in our cohort are members of Clalit health services who live in the southern part of Israel (the Negev), and receive most of their medical services at SUMC (located in Beer-Sheva). This population represents ~75% of the ~700,000 citizens in the Negev (Israel 2016), and is composed of ~60% Jewish and ~40% Bedouin Arabs: two ethnic groups that differ in their genetic background and lifestyle. The population of the Negev is relatively static such that many multigenerational families receive medical services at SUMC throughout their entire lives.

### Participant evaluation and recruitment

Children who are referred to the Child Development Institute (CDI) or to the Preschool Psychiatric Unit (PPU) at SUMC with suspected social communication difficulties and\or repetitive behaviors go through a rigorous clinical assessment that includes a comprehensive intake interview regarding the clinical and socio-demographic background of the diagnosed child, assessment with the Autism Diagnostic Observation Scale-2 (ADOS-2) (Lord et al. 2000) test, and a cognitive evaluation using either the Bayley Scales of Infant and Toddler Development- third edition (Bayley-III) (Bayley 2006) or the Wechsler Preschool and Primary Scale of Intelligence – version three (WPPSI-III) (Wechsler 1989) (Figure 1). Diagnosis of autism is determined by a child psychiatrist or a pediatric neurologist according to DSM5 criteria (American Psychiatric Association 2013). Parents or the legal guardian of all referred children are asked to enroll in the study and sign a consent form. This study was approved by the Helsinki committee responsible for human studies at SUMC.

Children with a positive diagnosis of autism are asked to return to the clinics for follow-up visits every 6 to 12 months until the age six. During these visits, their diagnosis is re-evaluated and families are invited to participate in additional ongoing experiments as detailed below.

**Figure 1:**
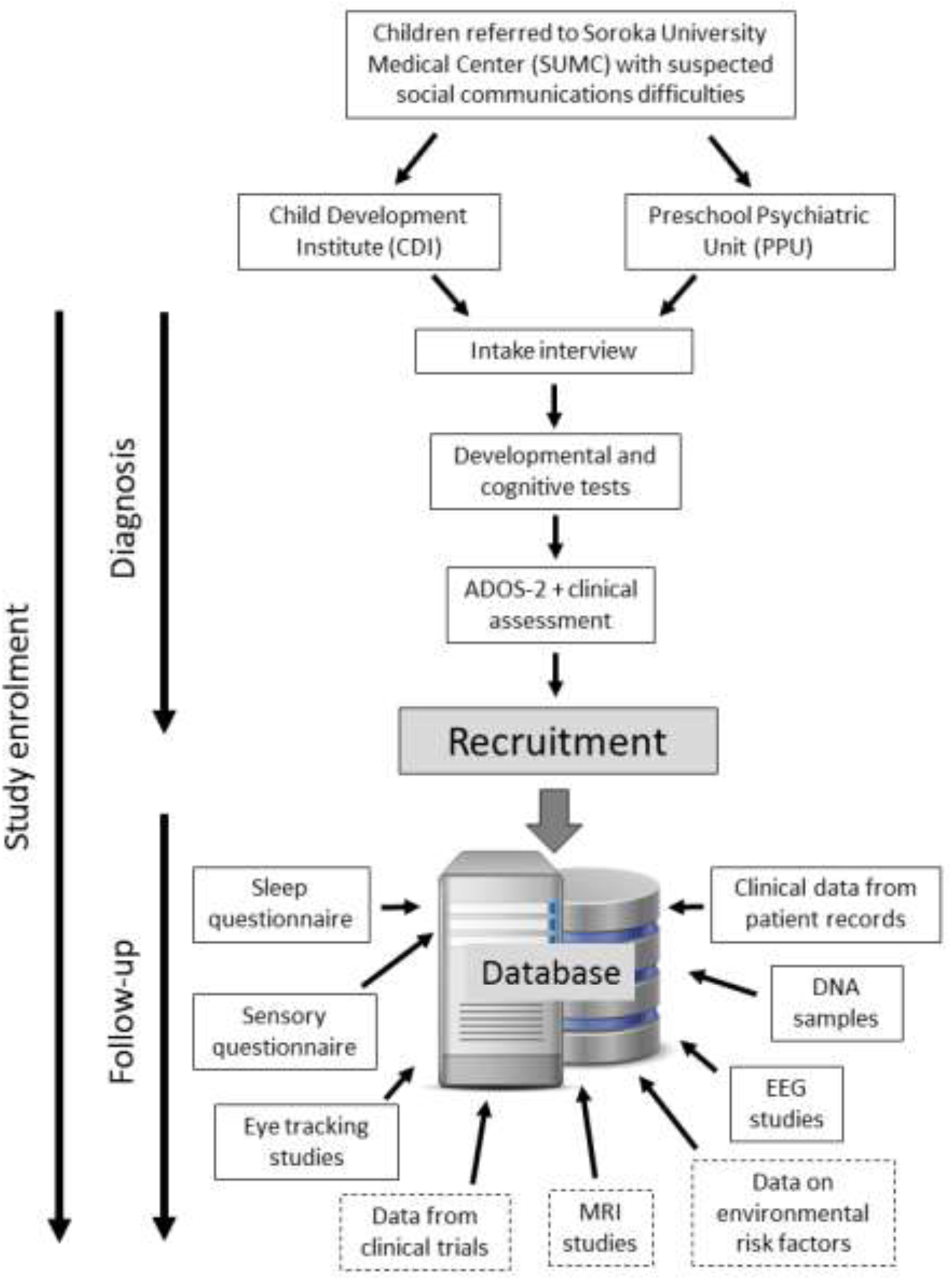
A flowchart of the evaluation of children and data collection processes in our cohort. Recruitment of families to the study is done either during the initial diagnosis, or during the follow-up visits that are also used to collect additional data. Currently collected data are presented in solid frames. Planned data collection is presented in dashed frames.

### Data collection and storage

Participating families are asked to complete the children’s sleep habits questionnaire (CSHQ, (Owens et al. 2000)) and the children’s sensory profile 2 questionnaire ((Dunn 2014)). In addition, we extract the existing clinical data of the children and their family members from the SUMC patient records. For example, we retrieve a range of potential prenatal and perinatal risk factors directly from the electronic database of the obstetrics and gynecology department (OGD) (the only operative maternity center in the Negev). This includes information about maternal characteristics, pregnancy and perinatal outcomes (e.g., gestational age, birth weight and Apgar scores), and peripartum maternal outcomes (e.g., hypertensive disorders, hypoxic-related complications, and gestational diabetes). The OGD database is used regularly for research purposes, and its accuracy is ensured through a standardized review of the data by a specialist medical secretary and a consulting obstetrician before it is coded (Amir et al. 2009).

Some of the recruited families are also enrolled in: 1) genetic studies where we extract DNA from a saliva sample of the affected child and their parents using Oragene-DNA kits (OG-575 or OG-500 for saliva samples). Aliquots of the extracted DNA are currently stored at −20^0^C and will be sent for whole genome sequencing. 2) an eye-tracking study where children are asked to watch movies in order to assess their ocular-motor control and social preferences. 3) a sleep-EEG and polysomnography study which is being performed at the SUMC sleep lab to better understand sleep architecture and early brain function in autism. We are currently expanding these data collection efforts so as to create a large overlap of measures across as many children as possible. All of these studies have also been approved by the SUMC Helsinki committee.

All of the data collected from participants in this cohort are organized and stored in a designated secured computerized database that has been certified by the Israeli ministry of justice according to the privacy protection law of Israel (Ministry of Justice 1981).

### Statistical analysis

We compared selected parameters between Jewish and Bedouin children using independent t-test or Mann-Whitney test for continuous variables, and Chi-square or Fisher-exact tests for nominal variables. We also used a 1-degree of freedom chi-square test for linearity to test for differences in trend of ordinal variables.

## Results

During the first eighteen months of the study (January 2015 to June 2016) 296 children (218 Jewish, 76 Bedouins, and 2 of mixed origin [Bedouin father and Jewish mother]) were referred to SUMC with suspected social communication difficulties (Figure 2). Autism diagnosis was confirmed in 188 of these children (63.5%), and 133 of these (70.4%) were successfully recruited to our study. Interestingly, the rate of positive autism diagnoses was significantly higher among Jewish children than among Bedouin children (68.3% vs 51.3%; P = *0.0077*). No significant differences in recruitment rates were found between the Jewish and Bedouins families.

**Figure 2:**
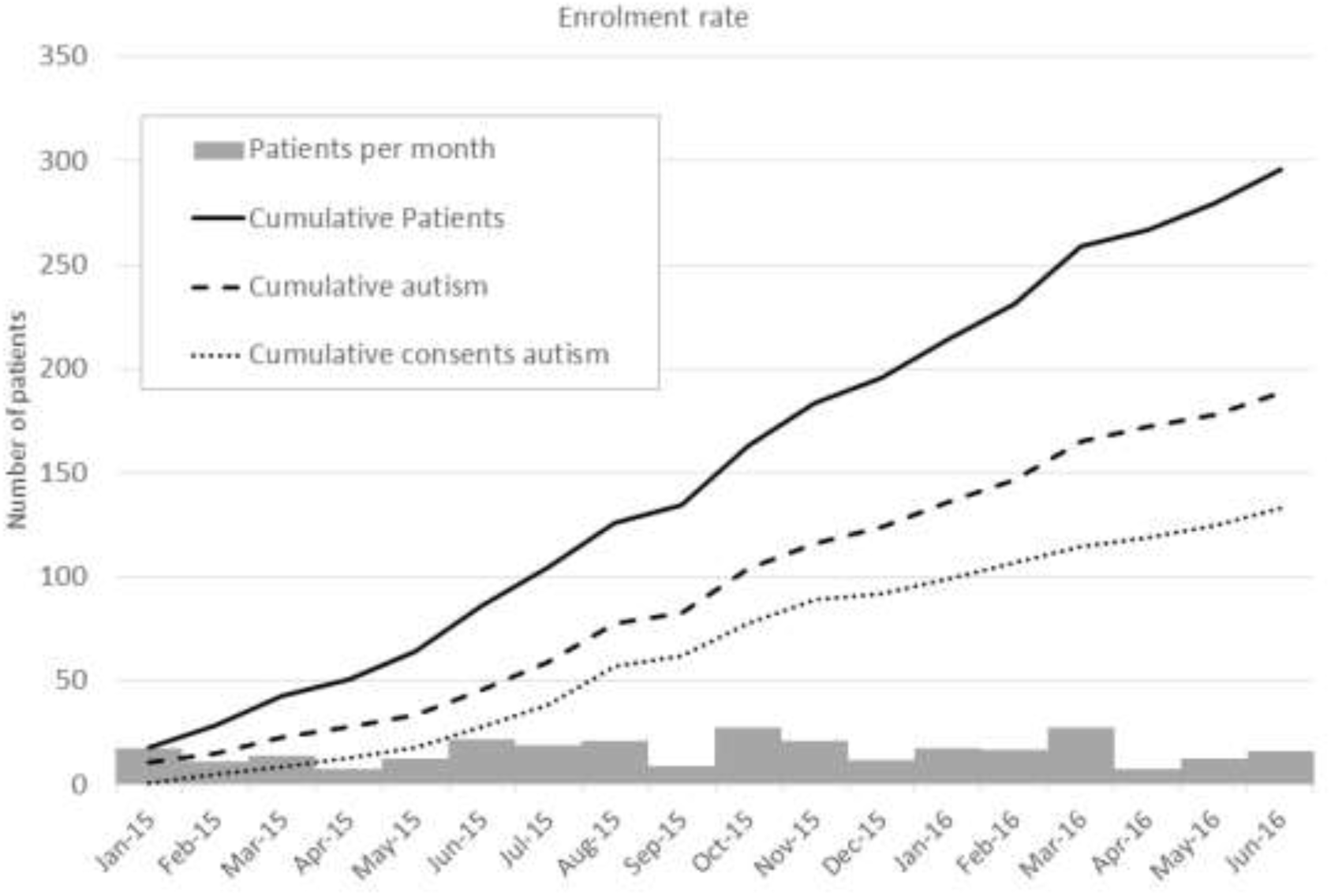
Growth of the Negev HUB database over an 18 month period. Families were referred to SUMC at an average rate of ~16.5 per month (range: 8-28 participants/moth; Gray bars). This resulted in a total of 296 participants who were evaluated during the first eighteen months of our study (continuous Black line). Of these, 188 children were diagnosed with autism (dashed Black line), and 133 of these signed the informed consent. Extrapolation of these data suggests that within 5 years of the study, we expect at least 444 children with autism and informed consent in our database.

Children diagnosed with autism included 154 males and 34 females (a male-to-female ratio of 4.5) who were 38.88 ± 15.82 months (range 16-98 months) at the time of diagnosis (Table 1). There was a higher proportion of females diagnosed with autism among Bedouins who were also 4.7-month younger at diagnosis compared to Jewish children (24.3% vs. 16.8%, and 35.20 vs. 39.91 respectively) however these differences were not statistically significant. We also didn’t find any significant differences in developmental milestones as reported by the parents (i.e. age at first crawl/walk/talk) between Bedouin and Jewish children with autism. Notably, the average maternal age at birth of Jewish mothers was five years older than that of Bedouin mothers (31.25 vs. 26.02; *P<0.0001*). Furthermore, Jewish mothers of children with autism were, on average, two years older than the general population of Jewish mothers at birth at SUMC in the last decade (31.25 vs. 29.22; *P<0.0001*). In contrast, Bedouin mothers of children with autism were, on average, slightly younger than the general population of Bedouin mothers at birth at SUMC in the last decade, but this difference was not statistically significant (26.02 vs. 27.15; *P = 0.367*). Jewish fathers of children with autism were, on average, slightly older at birth than Bedouin fathers of children with autism, but this difference was not statistically significant (33.40 vs. 31.01; *P = 0.135*).

All four modules of ADOS-2 were used for autism diagnosis in this sample. However, there were significant differences in module utilization in the diagnosis of Bedouin and Jewish children (*P<0.0001)*. Specifically, there was a higher tendency to utilize the 1^st^ module of the ADOS-2 (used with non-verbal children) over modules 2 or 3 among Bedouin children in comparison to Jewish children. Similarly, cognitive function was more commonly assessed with the BAYLYS test in Bedouin children as opposed to the WPPSI test, when compared with the Jewish children (*P=0.009*). These differences suggest a higher proportion of language difficulties among Bedouin children at the time of diagnosis in comparison to the Jewish children. In addition, Bedouin children exhibited significantly lower cognitive function compared to Jewish children (*P=0.013*), but there were no significant differences in the levels of autism severity between the two ethnic groups (see Table 1).

**Table 1:** Sample characteristics of children with autism at the Negev HUB database

| Variable | Bedouins | Jewish | Total | P-value |
| --- | --- | --- | --- | --- |
| Number of children <sup>a</sup> | 37 (20.7%) | 149 (79.3%) | 186 (100%) |  |
| Number of females | 9 (24.3%) | 25 (16.8%) | 34 (18.3%) | 0.280 <sup>c</sup> |
| Diagnosis Age (Months) <sup>b</sup> | 35.20 ± 12.69 | 39.91 ± 16.57 | 38.88 ± 15.82 | 0.152 <sup>d</sup> |
| Mother age at birth (Yeas) <sup>b</sup> | 26.02 ± 4.36 | 31.25 ± 5.21 | 30.27 ± 5.43 | < 0.0001 <sup>d</sup> |
| Father Age at birth (Years) <sup>b</sup> | 31.01 ± 8.63 | 33.40 ± 5.85 | 32.97 ± 6.43 | 0.135 <sup>d</sup> |
| <u>Developmental milestones</u> |  |  |  |  |
| Age at first crawl (Months)* | 8.75 ± 3.58 | 7.70 ± 3.30 | 7.93 (3.37 | 0.714 <sup>e</sup> |
| Age at first walk (Months)* | 16.95 ± 8.31 | 14.59 ± 3.44 | 15.03 ± 4.78 | 0.926 <sup>e</sup> |
| Age at first talk (Months)* | 14.70 ± 10.34 | 17.27 ± 12.79 | 16.90 ± 12.43 | 0.395 <sup>e</sup> |
| <u>ADOS module used</u> |  |  |  |  |
| 1 | 24 (64.9%) | 45 (30.2%) | 69 (37.1%) | < 0.0001 <sup>c</sup> |
| 2 | 0 (0%) | 45 (30.2%) | 45 (24.2%) |  |
| 3 | 0 (0%) | 11 (7.4%) | 11 (5.9%) |  |
| Toddlers | 13 (35.1%) | 48 (32.2%) | 61 (32.8%) |  |
| <u>ADOS score (modules 1-3)</u> |  |  |  |  |
| Low | 6 (25.0%) | 22 (21.8%) | 28 (22.4%) | 0.642 <sup>f</sup> |
| Moderate | 9 (37.5%) | 52 (51.5%) | 61 (48.8%) |  |
| High | 9 (37.5%) | 27 (26.7%) | 36 (28.8%) |  |
| <u>ADOS score (Toddlers)</u> |  |  |  |  |
| Mild-to-Moderate | 2 (15.4%) | 8 (16.7%) | 10 (16.1%) | 0.641 <sup>g</sup> |
| Moderate-to-Severe | 11 (84.6%) | 40 (83.3%) | 51 (83.9%) |  |
| <u>Cognitive test</u> |  |  |  |  |
| BAYLEY | 19 (90.5%) | 58 (62.4%) | 77 (67.5%) | 0.009 <sup>g</sup> |
| WPPSI | 2 (9.5%) | 35 (37.6%) | 37 (32.5%) |  |
| <u>Cognitive level</u> |  |  |  |  |
| Extremely Low | 10 (47.6%) | 20 (21.5%) | 30 (26.3%) | 0.013 <sup>f</sup> |
| Borderline | 4 (19.0%) | 21 (22.6%) | 25 (21.9 %) |  |
| Low-Average | 5 (23.8%) | 25 (26.9%) | 30 (26.3%) |  |
| Average-High | 2 (9.5%) | 27 (29.0%) | 29 (25.4%) |  |

<sup>a</sup> Two children with mixed Bedouin-Jewish origin were excluded from this table
<sup>b</sup> Mean ± SD
<sup>c</sup> Chi-square; <sup>d</sup> Independent t-test; <sup>e</sup> Mann-Whitney; <sup>f</sup> Chi-square for trend; <sup>g</sup> Fisher-exact test

Our sample has several other interesting characteristics: 1) Ten children in our cohort have siblings with autism (multiplex families). 2) there are two pairs of monozygotic twins of which both siblings were diagnosed with autism and another thirteen pairs of dizygotic twins of which only one of the children was diagnosed with autism. 3) thirteen Bedouin children in our sample (35.1% of the Bedouin cases) and one Jewish child were from consanguineous families (parents were first cousins). 4) forty Jewish families and twelve Bedouin families reported regression in their child’s development prior to the diagnosis of autism. 5) >90% of the children in our cohort were born at SUMC (all of the Bedouin and 88% of Jewish children), thereby enabling us to retrieve data from their birth clinical records to search for pre-and-perinatal risk factors.

## Discussion

The Negev HUB database contains a wide variety of socio-demographic, behavioral, and clinical measures from a cohort of children with autism who were diagnosed and continue to be followed-up at SUMC. An important aspect of this cohort is its unique ethnic composition that includes Jewish and Bedouin families, two populations that are known to differ in a number of clinical and demographic characteristics (Bilenko et al. 2014; Gotsman et al. 2015; Kridin et al. 2016; Lazarev et al. 2014; Leshem et al. 2015; Shental et al. 2010; Shimony et al. 2009; Smirnov et al. 2016; Treister-Goltzman et al. 2015). Indeed, we observed significant differences in maternal age at birth, language capabilities at time of diagnosis, and cognitive abilities at time of diagnosis in our preliminary analysis. Other differences are likely to be found in the data that we collect from these families in our complementary studies. The differences between Jewish and Bedouin children in our cohort demonstrate the common differences in behavioral and clinical characteristics found across different ethnic groups in other sites (e.g. (CDC 2014)). These differences may be a reflection of ethnic disparities in risk factors and actual etiologies of autism which may have a significant impact on the prevalence and severity of autism in different human subpopulations. It is therefore of paramount importance to understand the effect of different risk factors on the different manifestation of autism among different human populations. In this realm, the Negev HUB autism database is extremely useful for comparing different ethnicities that clearly differ in their clinical characteristics and are likely to also differ in their genetic, environmental, neuroimaging risk factors and actual etiologies.

The fact that >90% of the children diagnosed with autism in SUMC were also born at SUMC and continue to receive most of their clinical services at the same hospital is a major advantage of our research effort. Participating families allow us to access the clinical records of the children and their immediate family members to examine multiple types of data ranging from MRI scans and EEG exams to blood tests and clinical anamnesis. A notable example of this is the computerized database of Obstetrics and Gynecology Department (OGD) at SUMC that contains data on all newborn infants. This database has already been used successfully to identify preand-perinatal risk factors for a wide variety of conditions (e.g. (Kessous et al. 2013a; Kessous et al. 2013b; Lebel et al. 2012; Pariente et al. 2013; Ratzon et al. 2011)) and embodies an excellent resource for studying the effect of these factors on autism susceptibility. Data from other units of SUMC such as the Division of Pediatrics and the Department of Neurology will be used to search for clinical conditions that are enriched among children with autism and their immediate family members.

It is increasingly accepted that autism is composed of a variety of overlapping clinical conditions which differ in their phenotype manifestation, underlying causes, and possibly effective treatments (Lai et al. 2013; Lane et al. 2014; Lord and Jones 2012). Accurate classification of these distinct autism subtypes, which is likely to have important practical clinical implications will require collection of a wide verity of behavioral, clinical, and biological measures from large cohorts over a sufficient period of time. We envision that this ongoing project will continue to accumulate longitudinal data over an unlimited period of time. Extrapolation of the current autism diagnosis rate to a period of five-years suggests that by December 2019, the database will include data from over 620 families of children with autism. We expect that the actual number of families will be larger for the following reasons: 1) The birth rate in this region is continuously growing (Israel 2006-2016). 2) The incidence of autism is continuously rising (Raz et al. 2014). 3) We anticipate more diagnoses of Bedouin children (who currently account for >50% of the births at SUMC) as awareness for the disorder improves and services become more readily available. We have previously collaborated with the Israeli Ministry of Health to promote autism awareness in the Bedouin community, an effort that successfully increased the rate of autism diagnoses at SUMC from nearly 0% to ~20% over a period of six years. We anticipate that this rate will continue to rise in the next years such that the number of Bedouin children diagnosed with autism will reflect their true prevalence in the population.

To maximize our ability to capture comprehensive information from each family, we have integrated our research team into the clinic, where they meet the families already during their first diagnostic visit. Thus, data collection is carried throughout the diagnosis process and families are not requested to make a special effort to participate in the research. This situation facilitates parental consent of more than 70% that ensures a fair representation of the local Negev population. Our research team also collects longitudinal data in a prospective manner from participating families via the follow-up clinical assessments that are scheduled for every 6 to 12 months. Behavioral and clinical measures are, therefore, acquired from the children at the earliest time-point possible (during their initial diagnosis), and as part of their regular follow-up visits to the clinic. This will enable us to follow the progress of these children across several developmental time points and assess the effects of different interventions and environmental factors on their development.

Our Negev HUB joins other examples of relatively large autism research databases that have been built in the last several years (e.g. (Croen et al. 2012; Fischbach and Lord 2010; Payakachat et al. 2015; Siegel et al. 2015; Stoltenberg et al. 2010)). These efforts are critical for studying autism heterogeneity and identifying specific autism etiologies that will likely benefit from distinct interventions. We expect that the existence of such HUBs in multiple international sites containing populations with different genetic backgrounds and environmental exposures will be crucial for achieving these goals.

## Compliance with Ethical Standards

### Funding

This study was funded by the Israeli Science Foundation (grant number 527/15).

### Ethical approval

All procedures performed in studies involving human participants were in accordance with the ethical standards of the institutional and/or national research committee and with the 1964 Helsinki declaration and its later amendments or comparable ethical standards.

### Informed consent

Informed consent was obtained from all individual participants included in the study.

